# Fission yeast cells overproducing HSET/KIFC1 provides a useful tool for identification and evaluation of human kinesin-14 inhibitors

**DOI:** 10.1101/257436

**Authors:** Masashi Yukawa, Tomoaki Yamauchi, Ken-ichi Kimura, Takashi Toda

## Abstract

Many cancer cells contain more than two centrosomes, yet these cancer cells can form bipolar spindles and appear to proliferate normally, instead of committing lethal mitoses with multipolar spindles. It is shown that extra centrosomes are clustered into two pseudo-bipolar spindle poles, thereby escaping from multipolarity. Human kinesin-14 (HSET or KIFC1), a minus end-directed motor, plays a crucial role in centrosome clustering and as such, HSET is essential for cell viability only in cancer cells with supernumerary centrosomes, but not in non-transformed cells. Accordingly, HSET is deemed to be an efficient chemotherapeutic target to selectively kill cancer cells. Recently, three HSET inhibitors (AZ82, CW069 and SR31527) have been reported, but their specificity, efficacy and off-target cytotoxicity have not been evaluated rigorously. Here we show that these inhibitors on their own are cytotoxic to fission yeast, suggesting that they have other targets in vivo except for kinesin-14. Nonetheless, intriguingly, AZ82 can neutralize overproduced HSET and partially rescue its lethality. This methodology of protein overproduction in fission yeast provides a convenient, functional assay system by which to screen for not only selective human kinesin-14 inhibitors but also those against other molecules of interest.

## INTRODUCTION

Most animal cells have two centrosomes from which mitotic bipolar spindles assemble. This bipolarity is essential for equal segregation of genetic material, thereby ensuring genome stability. Like DNA duplication, a cell has a robust regulatory mechanism by which centrosome number is maintained strictly as one or two copies per cell, which orchestrates the chromosome cycle (Conduit et al., 2015; Fu et al., 2015). Interestingly, it is known that in many cancer cells, this synchrony between centrosome and chromosome cycles becomes uncoupled, by which such cells contain more than two centrosomes. Nonetheless, these cancer cells appear to divide normally by means of bipolar spindles without undergoing lethal multipolar mitoses (Quintyne et al., 2005). It has been shown that these cells could form pseudo-bipolar spindles by clustering the supernumerary centrosomes into two poles (Cosenza and Kramer, 2016; Gergely and Basto, 2008). Perturbations in centrosome clustering trigger multipolar spindle formation and mitotic catastrophe specifically in cancer cells with supernumerary centrosomes (Kwon et al., 2008; Quintyne et al., 2005).

Centrosome clustering is achieved by a side-by-side, rather than dispersed, positioning of individual centrosomes; this configuration results in the formation of two spindle poles that is facilitated through a microtubule-dependent inward force. Accordingly, a variety of cellular processes affecting microtubule-based tension and motility are involved in the clustering of supernumerary centrosomes (Godinho et al., 2009; Kramer et al., 2011; Leber et al., 2010; Rhys et al., 2018).

One of the crucial factors required for centrosome clustering is human HSET/KIFC1. This protein is a member of the kinesin-14 family of proteins that includes mouse Kifc2, *Xenopus* XCTK2, *Drosophila* Ncd, *C. elegans* KLP-15, *Arabidopsis* ATK5 and KCBP, *Aspergillus* KlpA, fission yeast Pkl1 and Klp2 and budding yeast Kar3 (Ambrose et al., 2005; Endow and Komma, 1996; Goshima and Vale, 2005; Hanlon et al., 1997; Meluh and Rose, 1990; O'Connell et al., 1993; Paluh et al., 2000; Robin et al., 2005; Saito et al., 1997; Troxell et al., 2001; Walczak et al., 1998; Yukawa et al., 2018). Kinesin-14 motor proteins have minus-end directionality and comprise three functional domains, an N-terminal tail domain, a central coiled-coil stalk domain and a C-terminal motor domain that possesses the ATPase activity (She and Yang, 2017). It is reported that HSET is abundantly expressed in several cancer cell lines including ovary, breast and lung cancer (Grinberg-Rashi et al., 2009; Pannu et al., 2015; Pawar et al., 2014). Intriguingly, knockdown of HSET in supernumerary centrosome-containing breast cancer cell lines prevents centrosome clustering and induces cell death by multipolarity in anaphase, while in normal cell lines that contain two centrosomes this treatment does not result in lethality (Kleylein-Sohn et al., 2012; Kwon et al., 2008). Therefore, specific targeting of HSET may provide a novel strategy by which to selectively kill cancer cells (Li et al., 2015; Xiao and Yang, 2016).

To date, three HSET inhibitors, designated AZ82, SR31527 and CW069, have been reported (Cosenza and Kramer, 2016; Xiao and Yang, 2016; Zhang et al., 2016) (Table 1). AZ82, which is the first HSET inhibitor to be identified, binds specifically to the HSET-microtubule binary complex, thereby inhibiting the microtubule-stimulated ATPase activity of HSET (Park et al., 2017; Wu et al., 2013; Yang et al., 2014). Upon addition to cancer cells with supernumerary centrosomes, this small molecule inhibitor triggers multipolar spindle formation and mitotic catastrophe. The second inhibitor, CW069, was designed and synthesized according to in silico computational modeling for HSET binding (Watts et al., 2013). This compound binds to HSET in an allosteric manner and reduces its ATPase activity in vitro. While cells treated with monastrol, a kinesin-5 inhibitor, exhibit mitotic arrest with monopolar spindles, co-treatment with CW069 suppresses monopolarity induced by monastrol. Finally, SR31527 was identified through a high-throughput screen based on an ATPase assay of HSET (Zhang et al., 2016). It inhibits HSET by binding directly to a novel allosteric site within the motor domain without involving microtubules. SR31527 reportedly prevents bipolar clustering of extra centrosomes in triple-negative breast cancer (TNBC) cells, and significantly reduces viability of TNBC cells. Despite the developments of these HSET inhibitors, their clinical efficacy has not been evaluated, nor has the possibility of offtarget cytotoxicity been scrutinized, which is a crucial step for clinical application of these drugs.

**Table 1:**
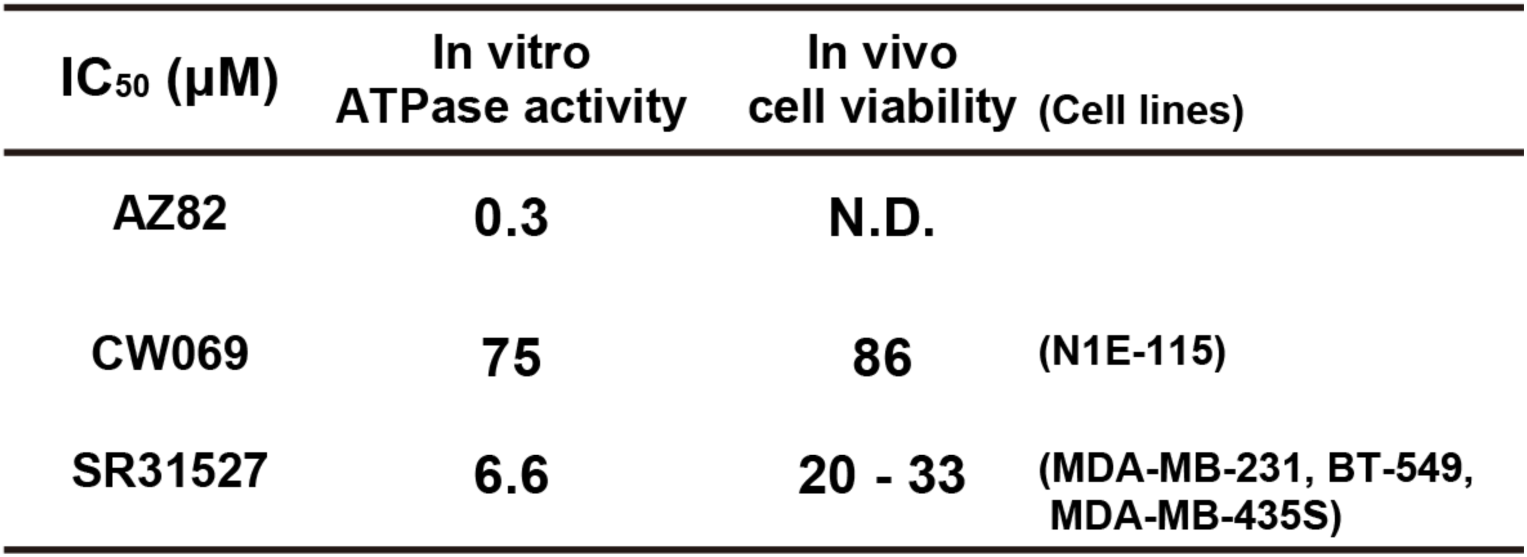
IC50 of HSET/KIFC1 inhibitors

Previously, we showed that HSET, when overexpressed in fission yeast, is toxic, and leads to mitotic arrest with monopolar spindles, reminiscent of overproduction of fission yeast kinesin-14, Pkl1 or Klp2 (Yukawa et al., 2018). In this study, we have attempted to exploit this phenotype for the functional evaluation of HSET inhibition.

We have found that all three reported HSET inhibitors have off-target effects on fission yeast growth. Intriguingly, however, AZ82 displays neutralizing activity against HSET-induced lethality.

## RESULTS

### The lethality of fission yeast cells overproducing HSET/kinesin-14 is rescued by co-overproduction of Cut7/kinesin-5

Bipolar spindle assembly requires proper force-balance generated by kinesin-5 and kinesin-14 (Sharp et al., 2000; She and Yang, 2017; Tanenbaum and Medema, 2010). In fission yeast, ectopic overproduction of kinesin-14 Pkl1 or inactivation of kinesin-5 Cut7 results in force imbalance leading to mitotic arrest with monopolar spindles (Hagan and Yanagida, 1990; Pidoux et al., 1996; Yukawa et al., 2018). Interestingly, mitotic arrest caused by Pkl1 overproduction is neutralized by co-overproduction of Cut7 and that cells overproducing both kinesins are capable of forming colonies (Rincon et al., 2017) (bottom two rows in Figure 1A).

**Figure 1:**
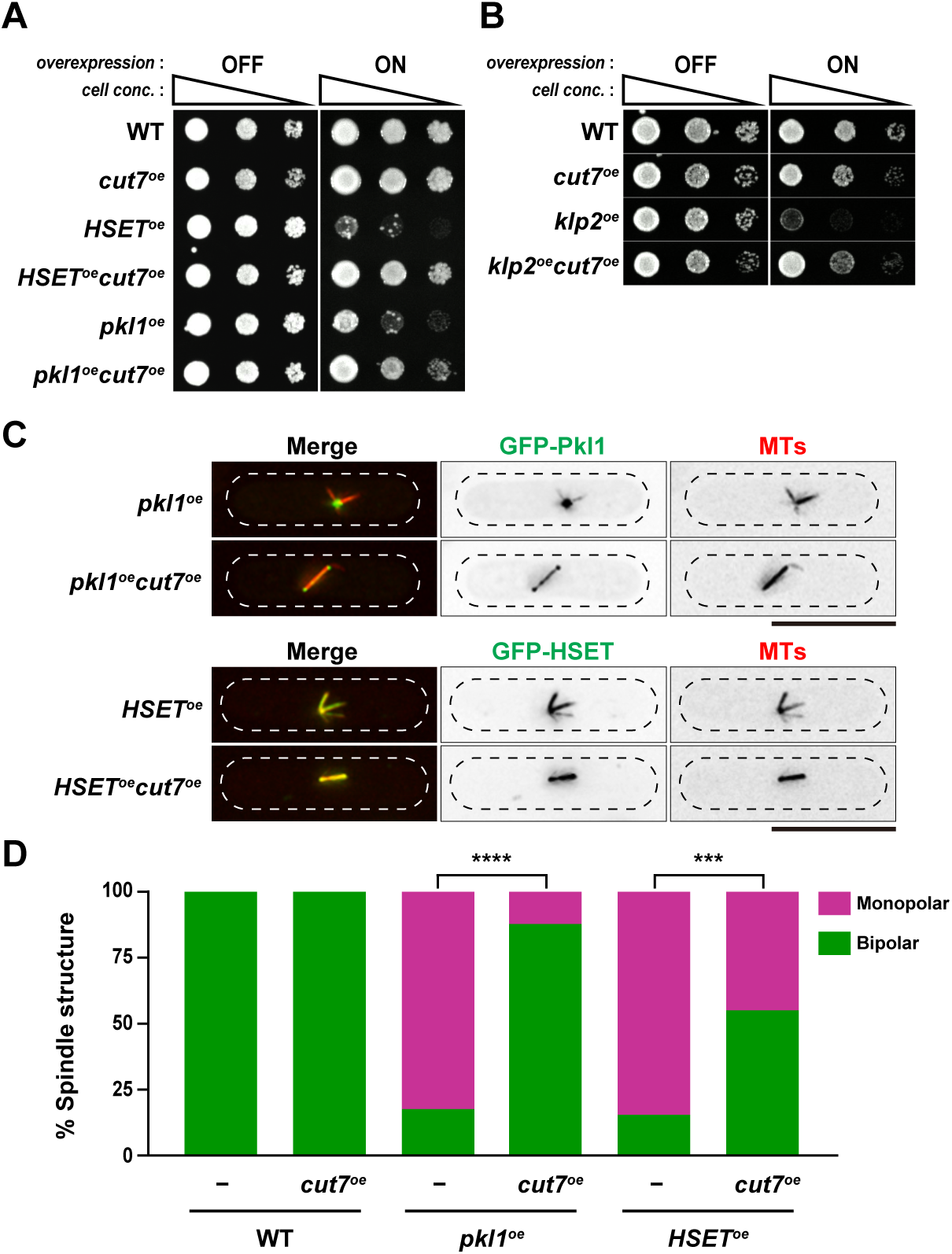
Co-overproduction of kinesin-5 Cut7 neutralizes growth toxicity derived from kinesin-14 overproduction. (**A, B**) Spot test. Strains overexpressing *cut7 (cut7^oe^)* and/or a gene encoding kinesin-14 (**A**, fission yeast *pkl1^oe^* or human *HSET^oe^* or **B**, fission yeast *klp2^oe^)* were serially (10fold) diluted, spotted onto minimal plates in the presence or absence of thiamine (+Thi/repressed or -Thi/derepressed, respectively) and incubated at 30°C for 3 d. *cell conc.,* cell concentration. *klp2* was expressed ectopically on plasmids. *nmtl-pkll* and *nmt41-HSET* were integrated at the *lysl* locus (Yukawa et al., 2018), while the *nmt41* promoter was integrated in front of the initiation codon of the *cut7* gene (Bahler et al., 1998). (**C**) Cellular localization of overexpressed Pkll and HSET and spindle morphology. Representative images are shown for each strain. Scale bars, 10 p,m. (**D**) The percentage of bipolar (green) or monopolar spindles (magenta) upon overexpression of either *pkll* or *HSET.* In each condition, more than 30 mitotic cells were counted (n>30). All P-values are derived from the two-tailed *%2* test (***P<0.001; ****P<0.0001).

We previously showed that overexpression of another kinesin-14, Klp2 or human HSET, is also lethal in fission yeast cells with a similar monopolar spindle phenotype (Yukawa et al., 2018). Therefore, we asked whether such Cut7-dependent rescue seen in Pkl1 overproducing cells is also observed in the case of overproduction of HSET or Klp2. As expected, Cut7 co-overproduction effectively suppresses the lethality of cells overproducing HSET or Klp2 (Figure 1A and B). Observation of spindle morphology showed that while cells overproducing only Pkl1 or HSET displayed a high frequency of monopolar spindles (84% or 85% respectively) (n=39), in those co-overproducing Pkl1 and Cut7 or HSET and Cut7, the frequency was substantially reduced to 12% or 45% respectively (n=38) (Figure 1C and D). Hence, HSET (and Klp2) is capable of generating inward pulling forces, thereby antagonizing outward pushing forces exerted by Cut7.

### An assay system for evaluation of human kinesin-14 inhibitors using fission yeast

Yeast-based screening for biologically active small molecules has successfully been implemented for the identification of promising drugs against human cancer and other diseases (Mager and Winderickx, 2005). This strategy is also useful to develop reagents that exhibit a beneficial impact on normal cells, e.g. those promoting lifespan extension (Sarnoski et al., 2017). Fission yeast has been used for several screenings, such as purification of specific compounds produced by *Actinomycetes* (Lewis et al., 2017) and a functional assay for Indinavir, an inhibitor against Human Immunodeficiency Virus Type-1 (Benko et al., 2016; Benko et al., 2017; Yang et al., 2012). Having seen toxicity derived from overproduced HSET in fission yeast cells (see Figure 1A), we exploited this lethal phenotype for the biological evaluation of known HSET inhibitors described earlier (AZ82, CW069 and SR31527) (Cosenza and Kramer, 2016; Xiao and Yang, 2016; Zhang et al., 2016).

Since we can detect the inhibitory activity of compounds through a simple assay for cell growth properties, it is possible to assess the specificity and the efficacy of these inhibitors. If inhibitors were truly specific to HSET molecules, these compounds on their own would not make any adverse impact on fission yeast growth (top row in Figure 2A, referred to as specific drugs). Upon overexpression of HSET, these inhibitors would now ameliorate viability loss resulting from HSET-mediated toxicity (bottom two rows in Figure 2A). By contrast, if these small molecules could not inhibit HSET activity, viability would not be increased in their presence (second row in Figure 2A). On the other hand, if inhibitors on their own interfere with fission yeast growth, it implies that these compounds recognize molecules or inactivate some pathways that are essential for fission yeast cells independent of HSET; in other words, these inhibitors are not specific to HSET, but instead they are drugs that exhibit off-target effects (top row in Figure 2B, referred to as multi-target drugs). Nonetheless, if HSET is effectively inhibited by these drugs, viability of fission yeast cells would be ameliorated to some extent compared to that without drug treatment (bottom row in Figure 2B).

**Figure 2:**
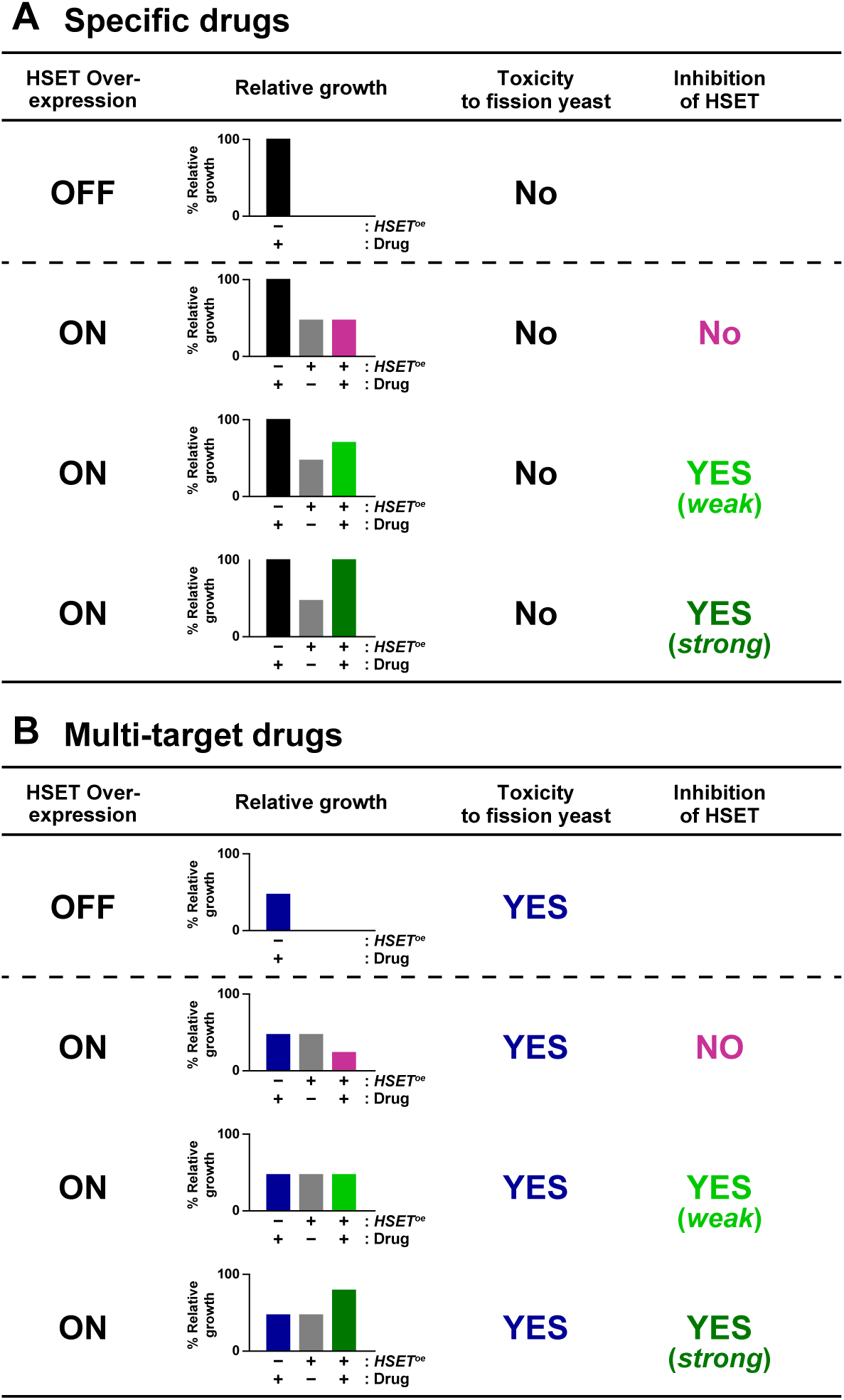
Strategy for evaluation of the specificity and efficacy of human kinesin-14 inhibitors using a fission yeast system. Inhibitors against human kinesin-14 are categorized into two classes. The first class is those that do not inhibit fission yeast cells on their own (**A**, specific drugs). The second class is those that possess growth-inhibiting activities on their own. It is likely that these drugs have some target molecules other than HSET that are essential for fission yeast cell viability (**B**, multi-target drugs). In either type A or B, we could assess inhibitory activity toward HSET by monitoring growth properties of cells in which HSET is overproduced in the absence or presence of drug treatment.

### All three known HSET inhibitors have non-specific cytotoxicity toward fission yeast cells

As aforementioned, if the HSET inhibitors were specific to human kinesin-14, addition of these molecules to wild type fission yeast should not incur any adverse effects on cell growth, as there are no targets (i.e. HSET) in these cells. Even if these inhibitors were to effectively inhibit fission yeast kinesin-14 molecules (i.e. Pkl1 and Klp2), they would not kill fission yeast cells, as deletion of either *pkl1* or klp2, or even double deletion is viable (Troxell et al., 2001). To address the specificity of HSET inhibitors, we first treated fission yeast cells individually with these drugs without introducing HSET. It is known that yeast cells are not highly sensitive to exogenous drugs for two reasons; not only does a thick cell wall prevent drugs entering the cells but the presence of P-glycoprotein transporters (multi-drug resistance proteins) actively pump drugs out of the cells (Nishi et al., 1992). Therefore, for our experiments, we exploited genetically tractable strains (YA8 and SAK931, Table 2) that are specifically designed for chemical biology (Arita et al., 2011; Takemoto et al., 2016b). In these strains, genes involved in influx and efflux of exogenously added drugs are multiply deleted; in YA8, genes encoding two major transporters, Bfr1 and Pmd1 are deleted *(bfrlΔpmdlΔ)* (Arita et al., 2011), while SAK931 contains deletions of 5 additional genes (7Δ) (Takemoto et al., 2016a).

**Table 2:** Fission yeast strains used in this study

| Strains | Genotypes | Figures used | Derivations |
| --- | --- | --- | --- |
| TY206 | *h<sup>-</sup> aur1R-Pnda3-mCherry-atb2 leu1 ura4* | 1A, 1D | This study |
| TY223 | *h<sup>+</sup> kanR-Pnmt41-cut7-kanR aur1R-Pnda3-mCherry-atb2 leu1 ura4 his2* | 1A, 1D | This study |
| MY1212 | *h<sup>-</sup> lys1<sup>+</sup>-Pnmt41-GFP-HSET aur1R-Pnda3-mCherry-atb2 leu1 ura4 lys1* | 1A, 1C-D | This study |
| TY241 | *h<sup>-</sup> kanR-Pnmt41-cut7-kanR lys1<sup>+</sup>-Pnmt41-GFP-HSET aur1R-Pnda3-mCherry-atb2 leu1 ura4 lys1* | 1A, 1C-D | This study |
| MY1230 | *h<sup>-</sup> lys1<sup>+</sup>-Pnmt1-GFP-pkl1 aur1R-Pnda3-mCherry-atb2 leu1 ura4 lys1* | 1A, 1C-D | This study |
| TY245 | *h<sup>-</sup> kanR-Pnmt41-cut7-kanR lys1<sup>+</sup>-Pnmt1-GFP-Pkl1 aur1R-Pnda3-mCherry-atb2 leu1 ura4 lys1* | 1A, 1C-D | This study |
| TY208 | *h<sup>-</sup> leu1 ura4 [pREP41-GFP]* | 1B | This study |
| TY212 | *h<sup>-</sup> kanR-Pnmt41-cut7-kanR leu1 ura4 [pREP41-GFP]* | 1B | This study |
| TY211 | *h<sup>-</sup> leu1 ura4 [pREP41-GFP-klp2]* | 1B | This study |
| TY215 | *h<sup>-</sup> kanR-Pnmt41-cut7-kanR leu1 ura4 [pREP41-GFP-klp2]* | 1B | This study |
| YA8 | *h<sup>+</sup> bfr1::ura4<sup>+</sup> pmd1::hisG leu1 ura4* | 3A-C | (Arita et al., 2011) |
| SAK931 | *h<sup>-</sup> caf5::bsdR pap1-Δ pmd1-Δ mfs1-Δ bfr1-Δ dnf2-Δ erg5::ura4<sup>+</sup> leu1 ura4 ade6-M210* | 3A | (Takemoto et al., 2016a) |
| MY1204 | *h<sup>-</sup> lys1<sup>+</sup>-Pnmt1-HSET bfr1::ura4<sup>+</sup> pmd1::hisG aur1R-Pnda3-mCherry-atb2 leu1 ura4 lys1* | 3B-C | This study |
| TY250 | *h<sup>-</sup> caf5::bsdR pap1-Δ pmd1-Δ mfs1-Δ bfr1-Δ dnf2-Δ erg5::ura4<sup>+</sup> leu1 ura4 ade6-M210 [pREP41-GFP]* | 3B-C | This study |
| TY247 | *h<sup>-</sup> caf5::bsdR pap1-Δ pmd1-Δ mfs1-Δ bfr1-Δ dnf2-Δ erg5::ura4<sup>+</sup> leu1 ura4 ade6-M210 [pREP41-GFP-HSET]* | 3B-C | This study |

As shown in Figure 3A, we found that all three drugs displayed very strong growth-inhibitory effects on YA8 or SA931 at 100 μM. CW069 was not cytotoxic in YA8 cells, but was in SAK931; fission yeast cells are likely less permeable to CW069 than AZ82 or SR31527. These results indicated that all three HSET inhibitors targeted molecules besides HSET within fission yeast cells that are essential for cell viability. It is noted that no obvious alterations in cell morphology were observed as a result of treatment with these drugs (Lewis et al., 2017), of which currently, the molecular details of off-target effects remain to be dissected. In summary we conclude that none of the three inhibitors examined is indisputably specific to HSET, but instead these small molecules interfere with unknown molecular pathways that are needed for cell viability of fission yeast.

**Figure 3:**
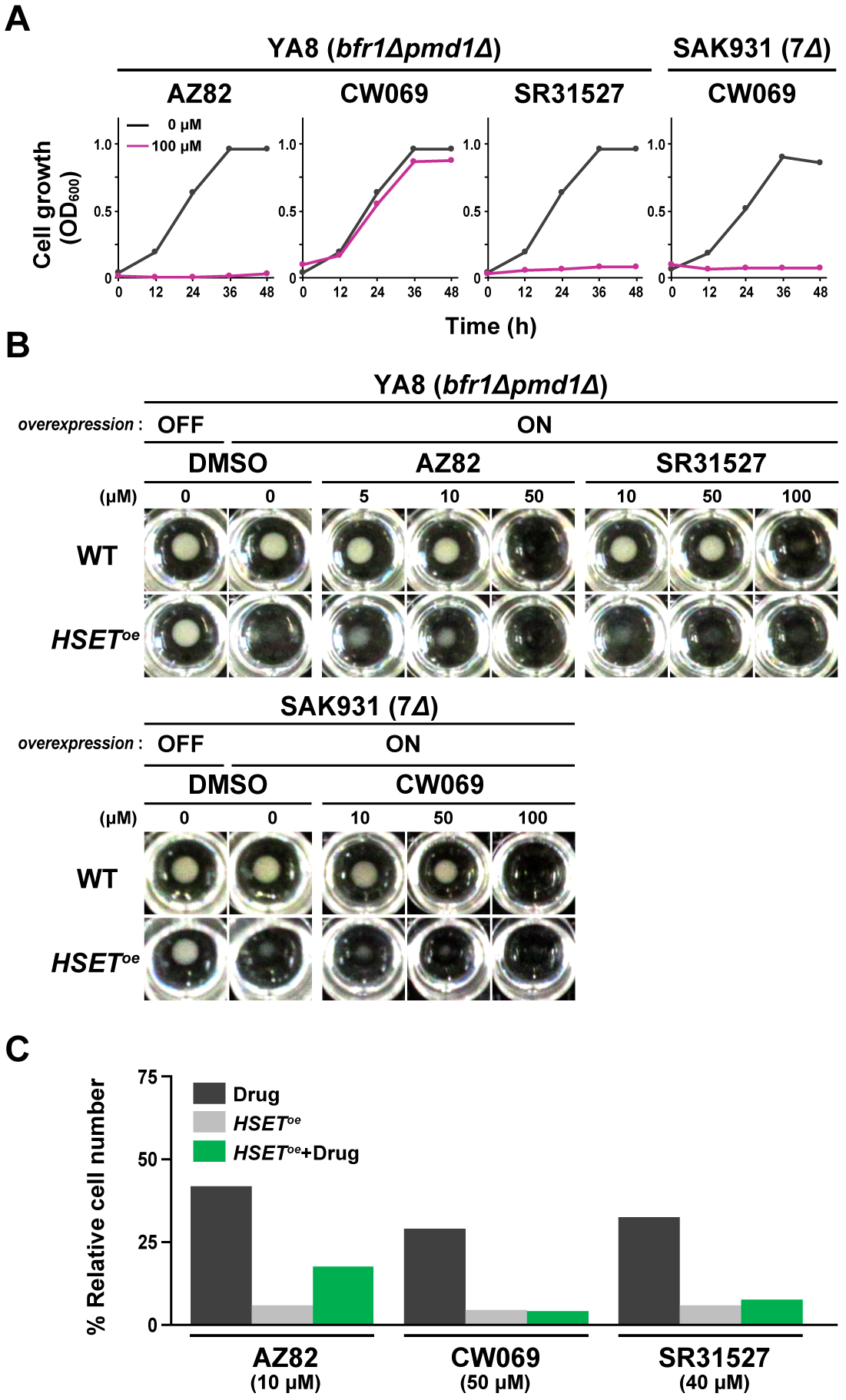
The three kinesin-14 inhibitors have off-target cytotoxicity in fission yeast, but AZ82 partially neutralizes HSET-induced lethality.

(**A**) Growth suppression of fission yeast cells upon treatment with individual kinesin-14 inhibitors. Two types of fission yeast strains (YA8 or SAK931) were used. (**B**) Growth characteristics of fission yeast cells upon HSET overproduction in the presence of individual inhibitors. YA8 or SA931 cells were inoculated at 10^5^ cells/ml in minimal medium in the absence of thiamine (derepressed condition) in 96-hole microplates, and DMSO (mock) or drug (dissolved in DMSO) was added. The final concentration of DMSO was 1%. Cells were incubated for 2-3 d at 30°C. WT: YA8 or SA931; *HSET^oe^* YA8 or SA931 overexpressing *HSET* gene. Note that small precipitates of cells are visible in *YA8/HSET^oe^* cells containing 5 or 10 μM of AZ82. (**C**) Relative number of HSET-overproducing cells in the presence of individual drugs. Cell number in the absence of drugs and *HSET^oe^* is set as 100%, and the relative value of individual conditions are calculated.

### AZ82 modestly rescues the lethality caused by HSET overproduction

We next sought to address whether these inhibitors are capable of rescuing the lethality of fission yeast cells resulting from forced expression of HSET. For this purpose, we assessed the number of cells in the presence or absence of inhibitors upon HSET overproduction. As the three kinesin-14 inhibitors displayed off-target cytotoxicity (Figure 3A), various concentrations of individual drugs were tested to find optimal concentrations which would, on one hand, repress HSET-mediated lethality, and yet, on the other hand display minimal cytotoxicity on their own. Intriguingly, we found that 10 μM AZ82 could rescue the lethality caused by HSET overproduction (~3 fold increase of viability; 6% vs 17% in the absence and presence of AZ82 respectively, Figure 3B and C). As the viability of cells treated with AZ82 (10 μM) alone dropped to 42%, the degree of rescue by AZ82 would be larger than 3 fold. By contrast, neither CW069 or SR31527 exhibited noticeable recovery of cell viability at any concentrations examined (Figure 3B and C). Nonetheless, it is noteworthy that no additive toxicity was observed between these drugs and HSET overproduction, inviting the possibility that these two drugs also somehow neutralized HSET-mediated lethality, albeit in a very modest manner. Taking these results together, we consider that the three small molecule HSET inhibitors can block HSET function in fission yeast, although they possess off-target effects (Table 3).

**Table 3:** Effects of HSET/KIFC1 inhibitors in fission yeast

|  | Toxicity to<br>*S. pombe* | Inhibition of<br>HSET | Rescue of<br>cell growth |
| --- | --- | --- | --- |
| AZ82 | YES | YES | YES |
| CW069 | YES | YES | NO |
| SR31527 | YES | YES | NO |

## DISCUSSION

In this study, we have introduced a novel, simple assay system using fission yeast that enables us to monitor the specificity and efficacy of known inhibitors against human kinesin-14 proteins. Recently, much attention has been attracted to mitotic kinesins as novel antitumor targets and several kinesin inhibitors including those used in this study have been developed as potential cancer therapeutics (Al-Obaidi et al., 2016; Cosenza and Kramer, 2016; Huszar et al., 2009; Ma et al., 2014; Ohashi et al., 2015; Xiao and Yang, 2016). Among them, human kinesin-14 inhibitors are regarded to be promising, as many cancer cells display dysregulation of the centrosome cycle, resulting in the emergence of supernumerary centrosomes. Yet, these cells escape from lethal multipolarity, which is attributed to HSET-dependent centrosome clustering (Cosenza and Kramer, 2016; Gergely and Basto, 2008; Kwon et al., 2008). These drugs are expected to have minimal impact on non-transformed cells, as HSET is not essential for normal mitosis in human beings. Consistent with this notion, all three HSET inhibitors (AZ82, CW069 and SR31527) have been shown to inhibit motor activities of HSET in vitro and display growth-suppressing characteristics to some extent in human cancer cells that contain supernumerary centrosomes (Watts et al., 2013; Wu et al., 2013; Zhang et al., 2016). However, whether these drugs are specific to HSET has not rigorously been evaluated. One reason for this pitfall is that depletion or inactivation of

HSET, a target of these drugs, in tumor cells results in low viability, which hampers accurate assessment of off-target effects in these cells.

Given this complication, the fission yeast system introduced in this work is robust. As fission yeast cells are devoid of HSET, the existence of off-target effects are easily monitored and reliably assessed. Our results indicate that all three drugs have undesirable targets in fission yeast that are essential for cell viability. Accordingly, we ponder that in principle, none of these reagents is ideal as a specific HSET inhibitor. It is of note that consistent with this proposition, the existence of target molecules other than HSET was previously pointed out for all three drugs (Watts et al., 2013; Wu et al., 2013; Zhang et al., 2016).

Despite off-target effects, we have found that AZ82 exhibits inhibitory activity toward otherwise toxic HSET overproduction in fission yeast and that the other two reagents, CW069 and SR31527, are likely to neutralize toxicity resulting from HSET overproduction, although their impact is less than that of AZ82. As we are able to monitor the specificity and the efficacy of HSET inhibitors by implementing this system, if new inhibitors were developed, our rapid and robust strategy would provide a powerful tool for their biological assessment in combination with a conventional human culture cell system.

In fission yeast, as in other organisms, simultaneous inactivation of kinesin-5 and kinesin-14 rescues lethality resulting from kinesin-5 inhibition alone (Civelekoglu-Scholey et al., 2010; Mountain et al., 1999; O'Connell et al., 1993; Pidoux et al., 1996;

Rincon et al., 2017; Rodriguez et al., 2008; Saunders et al., 1997; Troxell et al., 2001; Wang et al., 2015; Yukawa et al., 2017; Yukawa et al., 2018). Thus, effective inhibitors against fission yeast kinesin-14s (Pkl1 and Klp2) could be identified using *cut7* temperature-sensitive mutants, as previously proposed in the *Aspergillus nidulans* system (Wang et al., 2015). These inhibitors might be also effective to suppress HSET activities, as HSET and Pkl1/Klp2 are structurally conserved and HSET functionally replaces for Pkl1 or Klp2 when introduced into fission yeast (Yukawa et al., 2018).

Finally, we would like to point out that the methodology described in this study could be exploited as an assay system for inhibitors against any human proteins of interest. Provided that overproduction of those human proteins display some defective phenotypes including lethality, which is often the case (Benko et al., 2017; Matsuyama et al., 2006; Nkeze et al., 2015), small molecule libraries, natural products or extracts from any organism could be used to identify potential inhibitors (Figure 4). In fact, we have recently obtained small molecules derived from plants that specifically rescue lethality caused by HSET overproduction with very little cytotoxicity on their own (M.Y., T.T., and K.K., unpublished results). Detailed characterization of these molecules is in progress using both fission yeast and human cancer cells.

**Figure 4:**
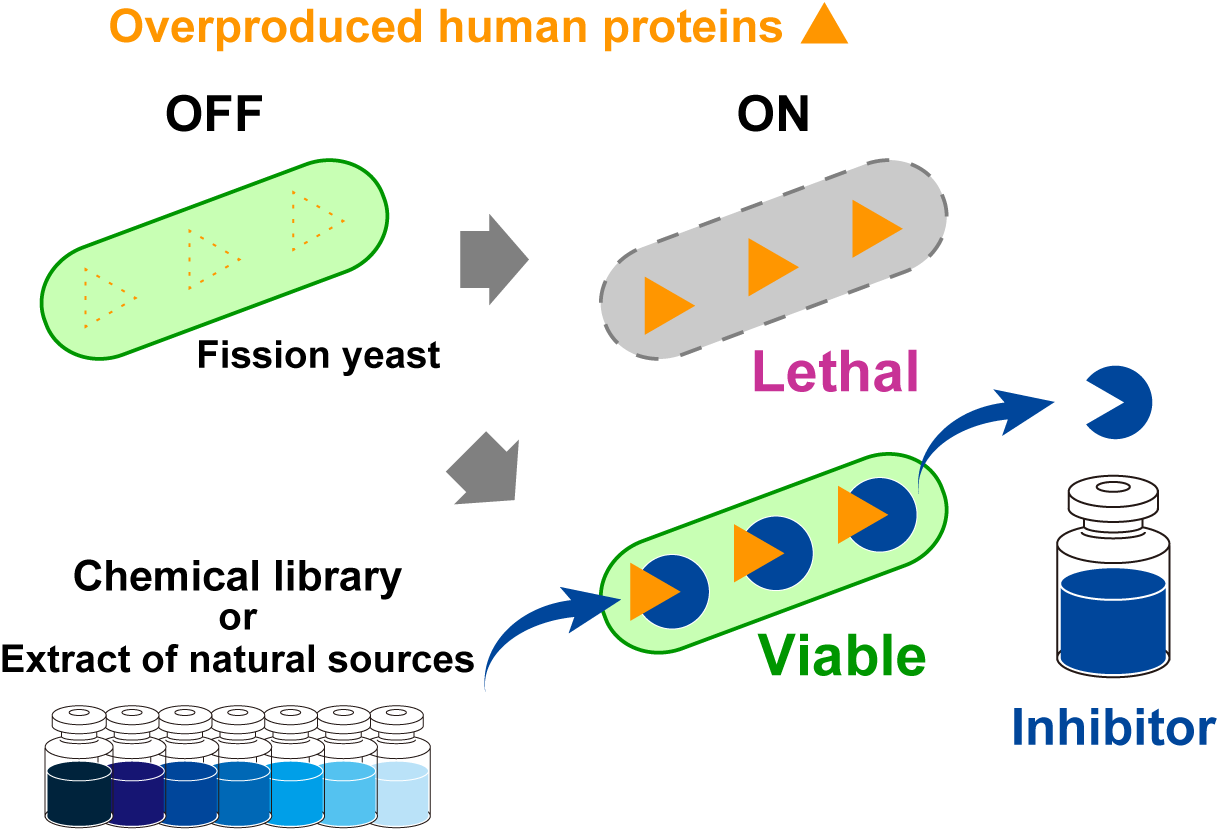
A general strategy for identification and evaluation of specific inhibitors against human proteins using a fission yeast system. Provided that human genes overexpressed in fission yeast confer some phenotype, such as lethality, this overproducing strain can be used as an assay system in which to identify specific inhibitors against these proteins. Libraries consisting of small molecules or natural products, or extracts prepared from *Actinomycetes,* marine organisms or plants can be used as sources of inhibitor compounds. Molecules that show activity that rescues lethal overproduction of protein X without any adverse offtarget effects toward fission yeast cells would be ideal.

## MATERIALS AND METHODS

### Strains, media, and genetic methods

Fission yeast strains used in this study are listed in Table 2. Media, growth conditions, and manipulations were carried out as previously described (Bähler et al., 1998; Moreno et al., 1991; Sato et al., 2005). For most of the experiments, rich YE5S liquid media and agar plates were used. Wild-type strain (513; Table 2), drug-sensitive strains (YA8 and SAK931) were provided by P. Nurse (The Francis Crick Institute, London, England, UK), M. Yoshida (Chemical Genetics Laboratory, RIKEN, Saitama, Japan) and S. A. Kawashima (Graduate School of Pharmaceutical Sciences, The University of Tokyo, Tokyo, Japan), respectively. For overexpression experiments using thiamine-repressible *nmt* series plasmids (Basi et al., 1993; Maundrell, 1990), cells were first grown in Pombe Glutamate Medium (PMG, the medium in which the ammonium of EMM2 is replaced with 20 mM glutamic acid) with required the amino acid supplements in the presence of 15 μM thiamine overnight. Thiamine was then washed out by filtration pump and cells continued to be cultured in the same PMG media in the absence of thiamine for further 12-24 h as necessary. Cells were serially diluted 10-fold, from an initial concentration of 2 × 10^7^ cells/ml, and spotted onto PMG plates with added supplements in the presence or absence of 15 μM thiamine. Plates were incubated at 27°C or 30°C.

### Preparation and manipulation of nucleic acids

Enzymes were used as recommended by the suppliers (New England Biolabs Inc. Ipswich, MA and Takara Bio Inc., Shiga, Japan).

### Chemical compounds

AZ82 was supplied from AstraZeneca (Boston, MA, U.S.A.). CW069 was a kind gift of S. V. Ley, the University of Cambridge. SR31527 was purchased from Vitas-M Laboratory (Apeldoorn, The Netherlands; Cat number: STK400735). All chemicals were dissolved in DMSO at 10 mM and stored at −20°C.

### Treatment of HSET-overproducing fission yeast cells with kinesin 14 inhibitors

Cells were cultured in YE5S or PMG liquid media with the required amino acid supplements and thiamine until mid-log phase at 30°C. Thiamine was washed out and cells shifted to the same PMG liquid media without thiamine at a concentration of 1 × 10^5^ cells/ml. The kinesin-14 inhibitors or DMSO (mock) were then added at various concentrations between 1 μM and 100 μM (0.01%-1% DMSO solution) and the cells were cultured at 30°C. OD measurements for yeast cultures were performed in a microplate reader (CHROMATE 4300; Awareness Technology Inc., Palm City, FL, U.S.A.), using an Iwaki Round Bottom 96-well microplate with lid and 200 μl per well for all measurements. OD measurements were performed at 600 nm at 30°C. Yeast cell number was determined using an automated cell counter (F-820; Sysmex, Kobe, Japan). Images of culture were taken by FAS-IV gel imaging system (Nippon Genetics, Tokyo, Japan).

### Fluorescence microscopy

Fluorescence microscopy images were obtained using a DeltaVision microscope system (DeltaVision Elite; GE Healthcare, Chicago, IL, U.S.A.) comprising a wide-field inverted epifluorescence microscope (IX71; Olympus, Tokyo, Japan) and a Plan Apochromat 6×, NA 1.42, oil immersion objective (PLAPON 60×O; Olympus Tokyo, Japan). DeltaVision image acquisition software (softWoRx 6.5.2; GE Healthcare, Chicago, IL) equipped with a charge-coupled device camera (CoolSNAP HQ2; Photometrics, Tucson, AZ, U.S.A.) was used. Live cells were imaged in a glass-bottomed culture dish (MatTek Corporation, Ashland, MA, U.S.A.) coated with soybean lectin and incubated at 27°C. To keep cultures at the proper temperature, a temperature-controlled chamber (Air Therm SMT; World Precision Instruments Inc., Sarasota, FL, U.S.A.) was used. Images were taken as 14-16 sections along the z axis at 0.2 μm intervals; they were then deconvolved and merged into a single projection. The sections of images acquired were compressed into a 2D projection using the DeltaVision maximum intensity algorithm. Deconvolution was applied before the 2D projection. Captured images were processed with Photoshop CS6 (version 13.0; Adobe, San Jose, CA, U.S.A.).

### Statistical data analysis

We used the two-tailed χ^2^ test to evaluate the significance of differences between frequencies of the cells with bipolar spindles in different strains. We used this key for asterisk placeholders to indicate p-values in the figures: e.g., ****, P < 0.0001.

## ACKNOWLEDGEMENTS

We are grateful to Fanni Gergely, Shigehiro A. Kawashima, Steven V. Ley, Paul Nurse, AstraZeneca (Xiaogang Pan), Yoko Yashiroda and Minoru Yoshida for providing us with strains and reagents used in this study. We thank Corinne Pinder and Tom Williams for critical reading of the manuscript and useful suggestions. We are grateful to Ayaka Inada for her technical assistance. This work was supported by the Japan Society for the Promotion of Science (JSPS) (KAKENHI Scientific Research (A) 16H02503 to T.T., a Challenging Exploratory Research grant 16K14672 to T.T., Scientific Research (C) 16K07694 to M.Y.), the Naito Foundation (T.T.) and the Uehara Memorial Foundation (T.T).

## AUTHOR CONTRIBUTIONS

M.Y. and T.T. designed research. M.Y. and T.Y. performed experiments and analyzed the data. M.Y. and T.T. wrote the manuscript, and T.Y., Y.K. and K.K. made suggestions.

